# Neural dynamics underlying self-control in the primate subthalamic nucleus

**DOI:** 10.1101/2022.10.24.513512

**Authors:** Benjamin Pasquereau, Robert S. Turner

**Affiliations:** Institut des Sciences Cognitives Marc Jeannerod, UMR 5229, Centre National de la Recherche Scientifique, 69675 Bron Cedex, France; Université Claude Bernard Lyon 1, 69100 Villeurbanne, France; Department of Neurobiology, Center for Neuroscience and The Center for the Neural Basis of Cognition, University of Pittsburgh, Pittsburgh, Pennsylvania 15261

## Abstract

The subthalamic nucleus (STN) is hypothesized to play a central role in neural processes that regulate self-control. Still uncertain, however, is how that brain structure participates in the dynamically evolving representation of value that underlies the ability to delay gratification and wait patiently for a gain. To address that gap in knowledge, we studied the spiking activity of neurons in the STN of monkeys during a task in which animals were required to remain motionless for varying periods of time in order to obtain food reward. At the single-neuron and population levels, we found a cost-benefit integration between the desirability of the expected reward and the imposed delay to reward delivery, with STN signals that dynamically combined both attributes of the reward to form a single integrated estimate of value. This neural representation of subjective value evolved dynamically across the waiting period that intervened between instruction cue and reward delivery. Moreover, this representation was gradually distributed along the antero-posterior axis of the STN such that the most posterior-placed neurons represented the temporal discounting of value most strongly. These findings highlight the selective involvement of the posterior STN in a dynamic internal process that dynamically estimates the ongoing cost-benefit balance of the current context. Such a process is essential for self-control, promoting goal pursuit and the willingness to bear the costs of time delays.

## INTRODUCTION

Imagine you are standing in a queue in front of a bakery. How long are you willing to wait for your favorite pastry? Many of us lose patience after about five minutes, while others persevere and keep waiting during longer delays. Our personal ability to delay gratification and maintain self-control depends on an internal process that estimates continuously the trade-off between the desirability of the benefit expected and the cost of waiting (Ainslie, 1975). All animals, including humans, prefer to receive rewards sooner rather than later, a phenomenon known as temporal discounting (Frederick et al., 2002; Loewenstein and Prelec, 1993; Mazur, 2001; Vanderveldt et al., 2016). Accordingly, people with low discount rates tend to pursue their long-term goals patiently, whereas people with high discount rates often abandon their goals impulsively and move on. In economic behavior, the net payoff for such a cost-benefit dilemma is typically evaluated by integrating the magnitude of the future reward with a hyperbolic discounting function (Green and Myerson, 2004; Kirby, 1997; Loewenstein et al., 1992). Most studies of the neural correlates of temporal discounting have focused on the task instruction period, the point in a trial when the subject is informed of the size of the reward to be delivered and of the delay in time until its delivery (Berns et al., 2007). Far less is known about neuronal activity during the subsequent post-instruction delay period, during which subjects may exhibit varying degrees of patience (e.g., self-control) and anticipation of reward. It is quite possible that neuronal representations evolve dynamically across this time period. For example, as the subjective value of a future reward is discounted by the passage of time, the motivation to achieve a delayed goal may decline gradually. Indeed, functional imaging studies in humans suggest that neural representations of temporal discounting evolve dynamically during a post-instruction delay period in patterns that differ distinctly between brain regions (Jimura et al., 2013; McGuire and Kable, 2015; Tanaka et al., 2020). How such dynamically evolving representations of temporally-discounted subjective value are instantiated at the single-unit level remains poorly understood.

Because the subthalamic nucleus (STN) is thought to be crucial in inhibitory control by preventing impulsivity (Aron et al., 2016; Bonnevie and Zaghloul, 2019; Jahanshahi et al., 2015) and modulating the performance of reward-seeking actions (Baunez et al., 2007; Baunez and Robbins, 1997), we hypothesized that this structure could contribute to the maintenance of adaptive behaviors by dynamically computing the temporally discounted value. The STN occupies a unique position for translating motivational drives into behavioral perseverance, standing at the crossroads between the basal ganglia indirect pathway and many prefrontal hyperdirect inputs involved in motivational, cognitive and motor functions (Haynes and Haber, 2013; Parent and Hazrati, 1995). Current functional models of the STN propose that increased activity in the STN extends the time to action initiation by elevating decision thresholds, preventing suboptimal early responses or decisions, especially in situations in which the motivational options are conflicting (Cavanagh et al., 2011; Frank, 2006; Mansfield et al., 2011). In support of these models, a series of lesion studies performed on rats has provided causal evidence that STN restrains premature responding in instrumental tasks (Baunez and Robbins, 1997; Wiener et al., 2008) and controls the willingness to work for food (Baunez et al., 2005, 2002). Dysfunctions of STN circuits even produced perseverative actions with a reduced ability to switch between behaviors (Baker and Ragozzino, 2014; Baunez et al., 2007), making this brain region a good candidate for regulating self-control and delayed gratification. Until now, however, existing evidence is mixed on whether the STN is causally involved in temporal discounting (Aiello et al., 2019; Evens et al., 2015; Seinstra et al., 2016; Seymour et al., 2016; Uslaner and Robinson, 2006; Voon et al., 2017; Winstanley et al., 2005), and no previous study has investigated how STN neurons process value information across delays.

Aside from its role in motor control, clinical studies support the involvement of the STN in motivational functions. In particular, deep brain stimulation (DBS-STN), which is effective at alleviating motor symptoms in parkinsonian patients, may induce a variety of side effects related to altered motivation such as depression, excessive eating behavior and hypomania (Berney et al., 2002; Castrioto et al., 2014; Jahanshahi et al., 2015; Voon et al., 2006). Electrophysiological recordings collected from the STN of these patients have shown low-frequency oscillations (<12Hz) and spiking activities related to various aspects of reward processing, with neural signals modulated by the magnitude of monetary reward and cost-benefit value attribution (Fumagalli et al., 2015; Justin Rossi et al., 2017; Zénon et al., 2016). In non-human animals, the ability of the STN to represent the subjective desirability of actions has also been evidenced by studies that show neurons firing as a function of the expected reward and the associated effort cost (Breysse et al., 2015; Espinosa-Parrilla et al., 2013; Nougaret et al., 2022). Although substantial effort has been directed to elucidate the role of the STN in valuation-related processes at the time of decision-making (Cavanagh et al., 2011; Coulthard et al., 2012; Frank et al., 2007) and in movement incentive (Nougaret et al., 2022; Tan et al., 2015; Zénon et al., 2016), much less attention has been paid to STN involvement in the computation of temporally discounted value during the post-instruction delay period, when behavioral inhibition must be sustained patiently over time. In addition, it is still unclear how these roles for STN in cost-benefit valuation and motivational processing relate to the known organization of this nucleus into anatomically and functionally distinct territories (Alexander et al., 1990; Nambu et al., 2002; Parent and Hazrati, 1995).

To determine if the STN conveys signals consistent with its predicted role in pursuing delayed gratification, we trained two monkeys to perform a delayed reward task in which animals were required to remain motionless during post-instruction delay periods of varying durations in order to obtain food reward. We hypothesized that STN neurons exhibit a dynamic encoding of temporally discounted value over the time course of the delay period consistent with a continuously evolving representation of value attribution essential for self-control. Here we tested this hypothesis by studying spiking activity in the STN while monkeys performed the task. At the single-neuron and population levels, our results support a role for the STN in temporal discounting, and indicate that neural signals underlying the valuation of reward size and delay are integrated dynamically into subjective value along an antero-posterior axis in this nucleus. Such dynamic value integration through the STN may regulate the expression of persistent behaviors for which a continuously evolving cost-benefit estimation is required to monitor and sustain goal achievement.

## RESULTS

### Monkeys’ behavior reflects reward size and delay in an integrated manner

Two monkeys (H and C) were trained to perform a delayed reward task in which they were required to align a cursor on a visual target and to maintain this arm posture for varying periods of time before delivery of food rewards (Fig. 1A). At the beginning of each trial, an instruction cue appeared transiently signaling one of six possible reward contingencies. Cue colors indicated the size of reward (1, 2, or 3 drops of food) and symbols indicated the delay-to-reward [short delay (1.8-2.3s) or long delay (3.5-4s)]. Animals were given the option to reject a proposed trial by moving the cursor outside of the target (e.g., if they did not think it was worth waiting for the expected quantity of reward). In this task, the rejection rate (i.e. the proportion of trials with a failure to keep the cursor in the target) reflects the monkey’s motivation to stay engaged in the task and to successfully complete the trial according to its prediction about the forthcoming reward. The six instruction cues effectively communicated six different levels of motivation or subjective value as evidenced by consistent effects on the animals’ task performance (Fig. 1B-C). Rejection rates were affected by both reward size (two-way ANOVAs; Monkey H: *F_(2,666)_*=10.47 *P*<0.001; Monkey C: *F_(2,708)_*=5.36 *P*=0.0049) and delay to reward (H: *F_(1,666)_*=22.62 *P*<0.001; C: *F_(1,708)_*=8.03 *P*=0.0047). Although the total proportion of rejected trials differed across monkeys (two-tailed t-test; *t*_(229)_=3.96 *P*<0.001), a similar behavioral pattern was observed between both animals during the task. The proportion of rejected trials was higher for smaller rewards and longer delays, while both animals waited more patiently to obtain larger rewards.

**Figure 1.**
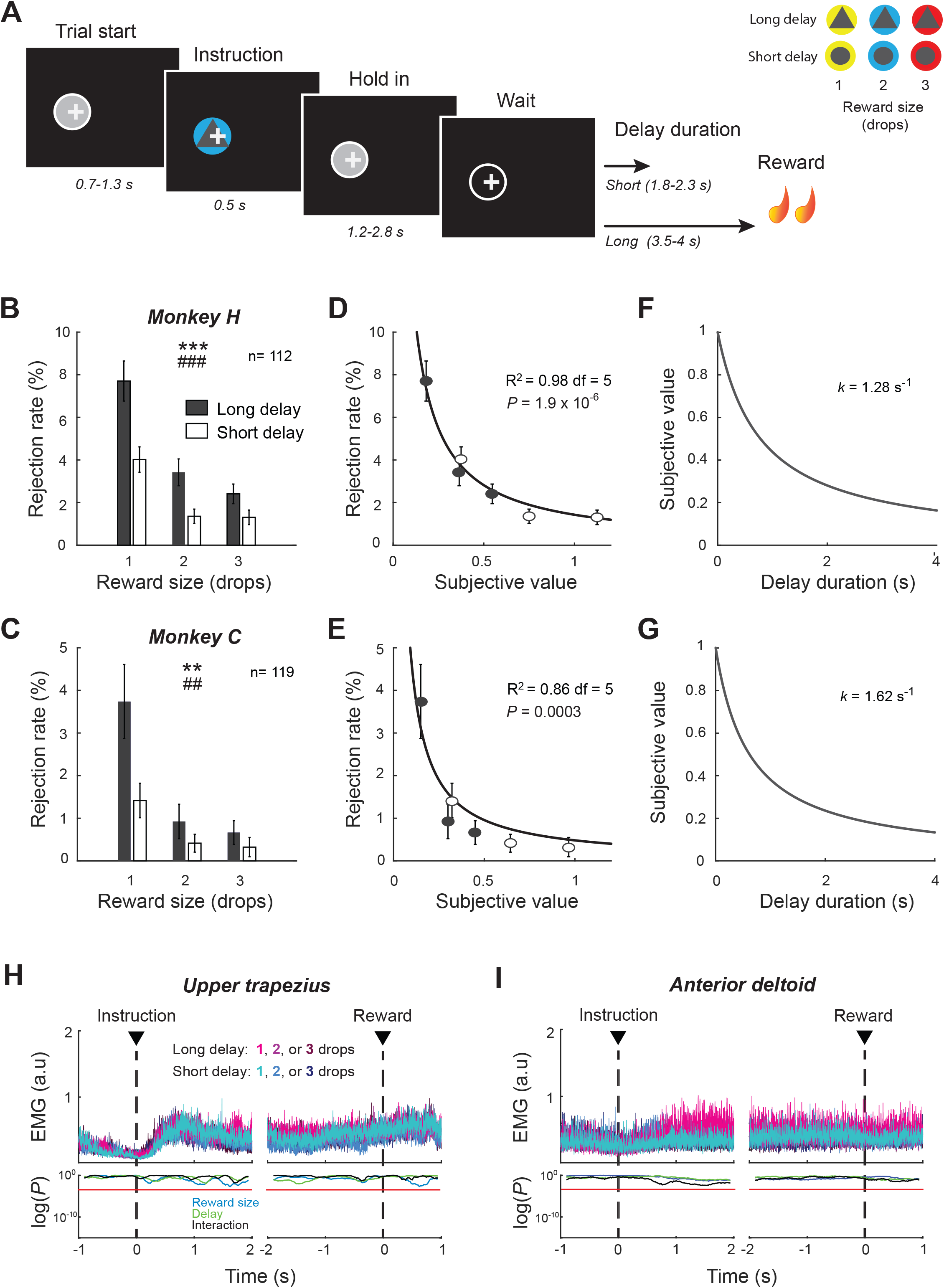
Delayed reward task and behavioral performance. (**A**) Temporal sequence of task events. After the monkey initiated a trial by positioning a cursor (+) within a visual target (gray circle), an instruction cue was presented briefly signaling the reward size (1, 2, or 3 drops of food) and the delayto-reward (short or long). The animal was required to maintain the cursor position over the waiting period to successfully obtain reward. (**B-C**) Rejection rates (mean ± SEM) were calculated and averaged for the six possible reward contingencies across sessions. Measures were affected by both reward size and delay (two-way ANOVA). Size: *** *P*<0.001 ** *P*<0.01; Delay: ### *P*<0.001 ## *P*<0.01. (**D-G**) For each animal, a temporal discount factor (*k*) was found that yielded the best fit between averaged rejections rates and the hyperbolic model expressed by Eq. 2. Goodness of fit was evaluated by the coefficient of determination (R^2^). (**H-I**) EMG signals collected in monkey H were aligned on the presentation of cues and reward. The effects of reward size and delay were examined using a series of two-way ANOVAs. Red lines indicate the statistical threshold (*P*<0.05 corrected for 143-time bins).

Interactive effects between reward size and delay (H: *F_(2,666)_*=19.31 *P*<0.001; C: *F_(2,708)_*=10.31 *P*<0.001) revealed an integration of both task parameters to estimate the overall desirability or subjective value of each cost-benefit condition. To characterize how subjective value declined with delay, we fitted the averaged rejection rates to a hyperbolic discounting model (as expressed by Eq. 2). To be more specific, we inferred the temporal discount factor (*k*) that maximized the inverse relation between each monkey’s behavior and the subjective value calculated from a hyperbolic function. Consistent with other monkey studies (Hori et al., 2021; Minamimoto et al., 2009), the animals’ task performance was well approximated by an inverse relation with hyperbolic delay discounting (H: R^2^=0.98; C: R^2^=0.86; Figs 1D-E). The resulting discount rates calculated for both animals were relatively close values (Figs 1F-G). In comparison, however, Monkey C was a bit more impatient with a steeper delay discounting (*k*=1.62 s^−1^), while the subjective value estimated by monkey H was slightly less impacted by the cost of waiting (*k*=1.28 s^−1^).

### Controls

We recorded EMGs from different muscles (trapezius, deltoid, pectoralis, triceps, biceps) while monkey H performed the behavioral task. During the post-instruction waiting interval, when the animal remained static, the maintenance of the arm posture resulted in a slight increase in the tonic activity of shoulder muscles (Figs 1H-I). As evidenced by a series of two-way ANOVAs (reward x delay, *P*<0.05 corrected for 143-time bins), muscle patterns were not altered by reward contingencies. This suggests that monkeys controlled their posture with a constant motor output across trial conditions, independent of reward size and delay. Alternatively, as monkeys were not required to control their gaze while performing the task, we found that their eye positions varied according to the type of trial (Fig.1-figure supplement 1). Eye positions were affected by both the expected reward size and delay after the presentation of instruction cues (two-way ANOVAs, *P*<0.05 corrected for 143-time bins). Reward-by-delay interactions detected in eye positions after instruction offset reinforce the view that cost-benefit parameters were integrated into a common valuation by monkeys.

To confirm the ability of our animals to recognize and evaluate appropriately the different instruction cues, the animals also performed a variant of the task that required decision making. In this variant, the monkey was allowed to choose freely between two alternate reward size/delay combinations. We observed appropriately strong preferences for the cues that predicted large rewards and short delays. Monkeys selected the more advantageous option in terms of reward when the delays were equal (H: 97%, *t*_(14)_=26.99 *P*<0.001; C: 99%, *t*_(16)_=30.55 *P*<0.001) and the more advantageous delay option when reward sizes were held constant (H: 99%, *t*_(11)_=29.71 *P*<0.001; C: 95%, *t*_(10)_=24.82 *P*<0.001).

### Neuronal activity of STN reflects reward size and delay

While the monkeys performed the delayed reward task, we recorded single-unit activity from 231 neurons in the right STN (112 from monkey H; 119 from monkey C). Similar to our previous study (Pasquereau and Turner, 2017), STN neurons were identified based on location and standard electrophysiological criteria (Figs 2A-C). Most STN neurons exhibited changes in firing rate at one or more times in the task. Approximately 37% of neurons demonstrated a peak in activity in the first second following presentation of the instruction cues, while 20% of neurons exhibited highest discharge rates just before reward delivery (Fig. 2D). Despite fact that phasic changes evoked by instruction cues dominated the population-averaged activity (Fig. 2E), we found that the variability of neuronal activities across the six reward/delay conditions was maintained at an elevated constant level across several seconds of the trial, as evidenced by the Fano Factor shown in Figure 2F. This suggests that task-relevant information was processed by STN neurons not only immediately after the presentation of the instruction cue, but also later in the course of the post-instruction delay period, when the animal was maintaining a stable arm position in anticipation of reward delivery.

**Figure 2.**
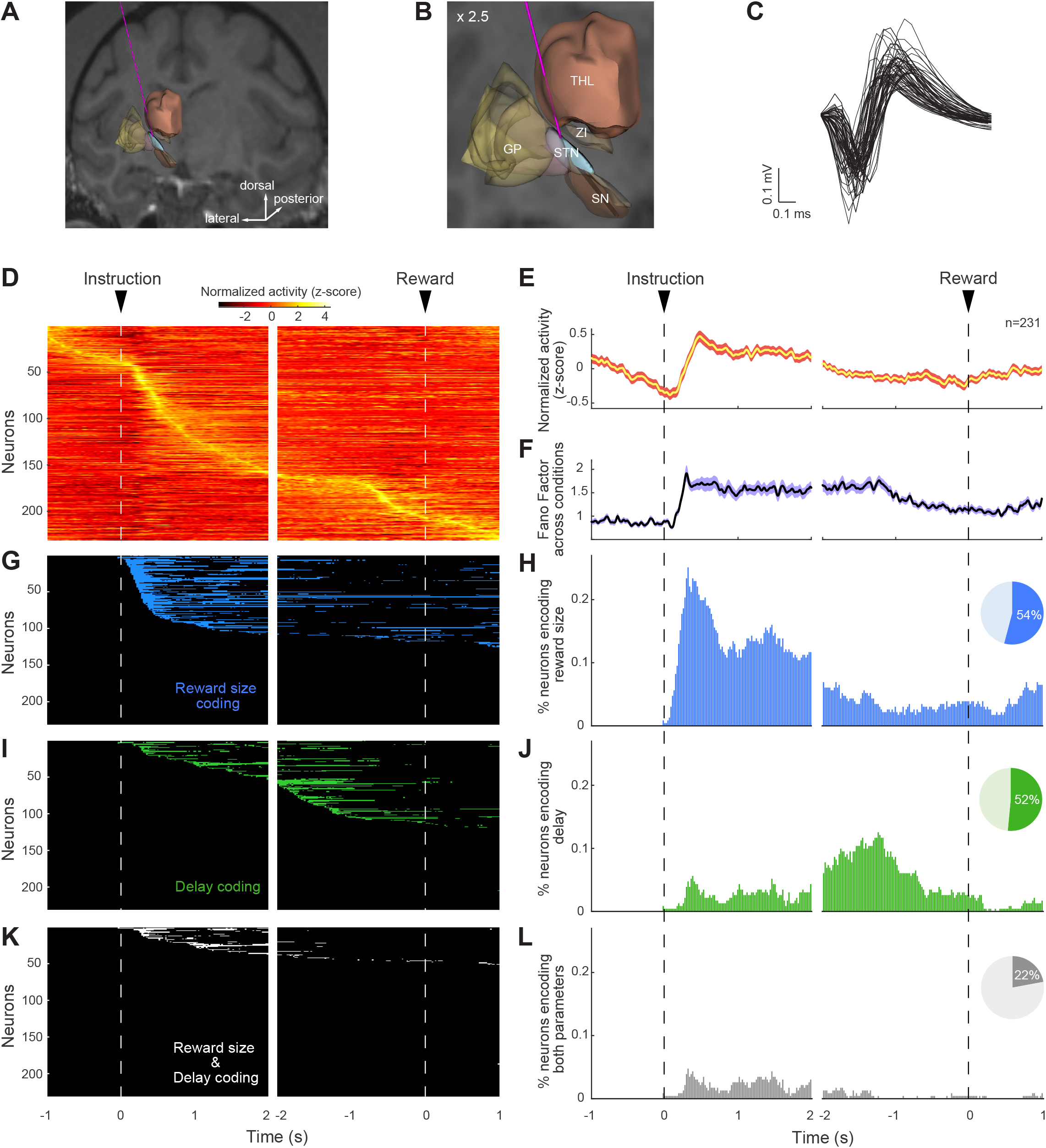
STN neurons were modulated by reward size and delay. (**A-B**) Reconstruction of a trajectory used for STN recordings with a structural MRI and high resolution 3-D templates of individual nuclei derived from (Martin and Bowden, 1996)). Globus pallidus (GP), Substantia nigra (SN), Zona incerta (ZI), Thalamus (THL). (**C**) Sample of action potential waveforms emitted by STN neurons. (**D**) Color map histograms of neuronal activities recorded from the STN. Each horizontal line indicates neural activity aligned to successive task events averaged across trial types. Neuronal firing rates were Z-score normalized. (**E**) Population-averaged activities of STN neurons, and (**F**) Fano factors that showed the variability of the neural population ensemble across the six possible reward contingencies. The width of the curves indicates the population SEM. (**G-L**) Influence of reward size and delay on individual neural activities was detected by a series of two-way ANOVAs (*P*<0.05, corrected for 188-time bins). The time course of encoding of task-relevant information (*left column*) and the fractions of neurons modulated by reward size and/or delay (*right column*) were represented for each time bin from instruction cue to reward delivery. Pie charts show the total fraction of STN neurons influenced by reward size (*blue*), delay (*green*), or both task parameters simultaneously (*gray*).

To determine whether and when STN neurons were involved in the evaluation of different task conditions, we tested the neural activities for effects of reward size and delay using two-way ANOVAs combined with a sliding window procedure (*P*<0.05, corrected for 188-time bins). Because the variability of neuronal activities across task conditions was sustained over time, we analyzed the spiking activity of each neuron in a time-resolved way across the course of trials from the presentation of the instruction cue to the onset of the reward. Of the 231 neurons recorded, 124 (54%) and 120 (52%) were modulated by reward size and delay, respectively (Figs. 2G-J). Interestingly, the two types of encoding occurred preferentially during different periods of the trial. Immediately following cue presentation, neurons were strongly influenced by the reward size signaled by the instruction while, later in the trial, as animals endured the waiting period, encoding of delay became more common. Among the 88 neurons (38%) sensitive to both parameters at some point over the course of the trial, 51 (22%) were simultaneously influenced by reward size and delay within the same time bins thereby enabling a direct integration of cost-benefit conditions by individual neurons (Figs. 2K-L). Reward-by-delay interactions were detected at a range of times across several seconds following the presentation of the instruction cue, suggesting that the way STN neurons represented task conditions changed dynamically across the course of a trial.

### Dynamic encodings of reward size and delay

To examine how reward size and delay were represented by individual STN units and how that representation changed across time in a trial, we performed time-resolved linear regressions with single-unit neural activity as the dependent variable. For each task-related neuron (i.e., neurons encoding at least one task parameter for a least one-time bin, n=156), we tested whether the firing rate was modulated by the expected reward quantity and the delay to reward delivery (as expressed by Eq. 3). Because the STN contains an oculomotor territory (Matsumura et al., 1992), we included measures of eye movements (i.e., gaze position and gaze velocity) in our model as nuisance variables. As illustrated in Figure 3 with three example neurons, task parameters were encoded in the STN at different stages of the trial following different modalities (2-tailed *t*-test, *P*<0.05). Based on the polarity of the regression coefficients β_*Reward*_, we found neurons whose activity transiently indexed reward size by increasing (e.g., neuron #1) or decreasing (e.g., neuron #2) their firing rate. Similarly, by detecting changes in the regression coefficients β_*Delay*_, we found neurons that increased (e.g., neuron #2) or decreased (e.g., neuron #1) their activity as a function of the delay to reward. Neural activities were often influenced in opposite directions by the predicted amount of reward and the delay (positive β_*Reward*_ with negative β_*Delay*_, or vice versa). The specific pattern of task encoding within individual cells, however, often changed over the course of the trial. For example, in the third exemplar unit activity shown in Figure 6 (right column) the influence of reward size on firing rate (i.e., β_*Reward*_) reversed repeatedly in the post-instruction epoch. This type of variability in the regression coefficients impeded simple approaches for categorization of STN neurons via their pattern of encodings (e.g., positive reward encoding vs. negative).

**Figure 3.**
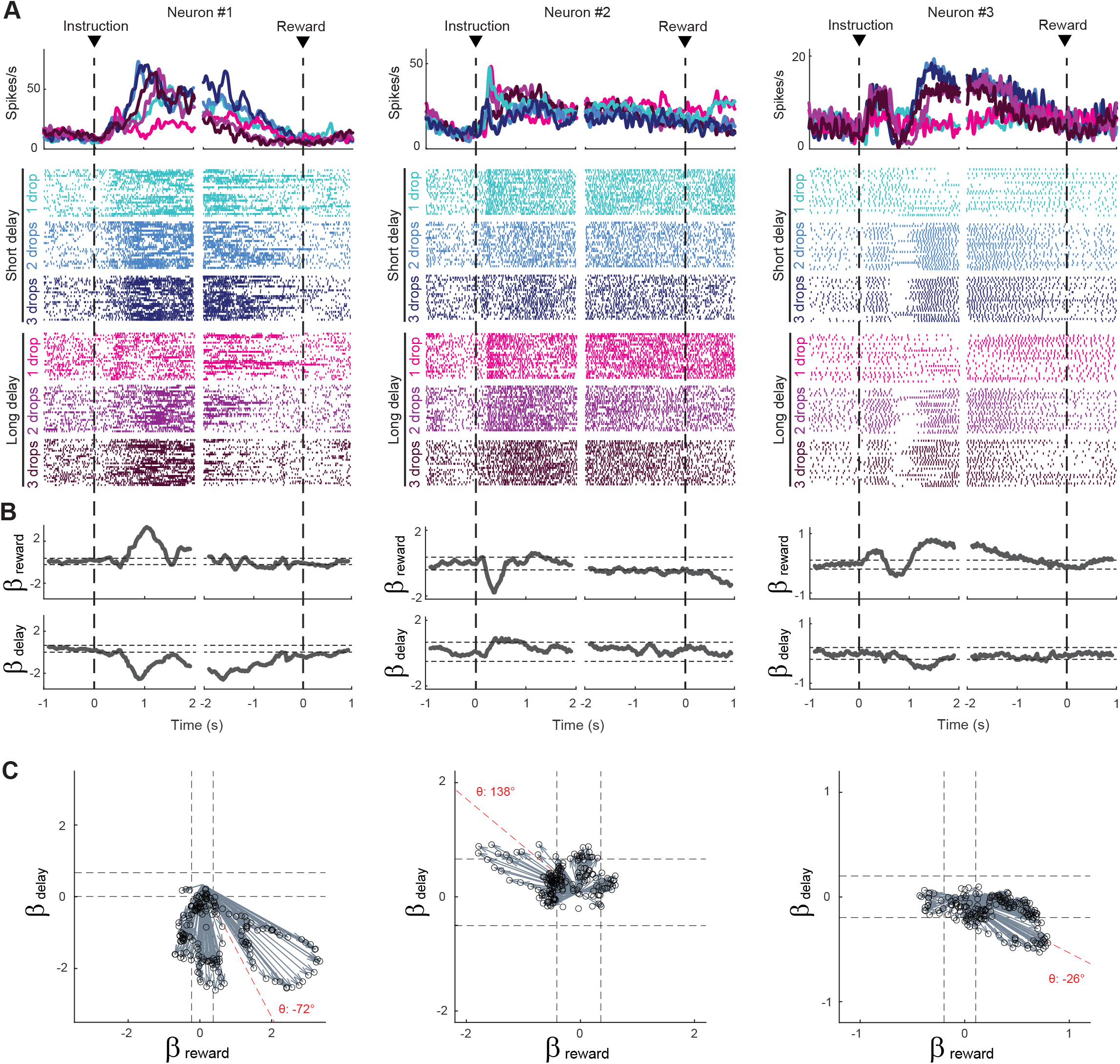
Response of STN neurons to the six possible reward contingencies. (**A**) The activity of three exemplar neurons that were classified as task-related cells. Spike density functions and raster plots were constructed separately around the presentation of instruction cues and the reward delivery for the different cost-benefit conditions. (**B**) A sliding window regression analysis compared firing rates between trial types (as expressed by Eq. 3). The regression coefficients (gray lines) were used to characterize the dynamic encoding of reward size (β_Reward_) and delay (β_Delay_). The horizontal dashed lines indicate the statistical threshold (2-tailed *t*-test, *P*<0.05). (**C**) Time series of regression coefficients projected into an orthogonal space where reward size and delay composed the two dimensions. Vector time series were produced for significant β values. Black dashed lines indicate statistical thresholds (*P*<0.05). The angle (θ) of the vector sum (*red dashed line*) was calculated to identify how neurons integrated cost-benefit conditions.

As a control, we evaluated the relative efficiency of our regression model, which disentangles reward size and delay (as expressed by Eq. 3), by comparing its goodness-of-fit against an alternate model in which neural activity is predicted by the subjective value estimated from an animal’s behavior (as expressed by Eq. 4, see Figs. 1D-E for details about subjective values). By comparing the values of the Akaike information criterion (AIC, a measure of goodness-of-fit) calculated for all recorded neurons, we found that our model (Eq. 3) provided a better fit to the population neural activity than the alternate model (AIC_model 3_ = 173.88 ± 7.36, AIC_model 4_ = 174.46± 7.38; paired *t*-test, *t*_(230)_=3.96 *P*<0.001).

### Mixed encodings of reward size and delay

To gain deeper insight into how the reward and delay dimensions of the task are integrated by a neuron’s activity, we projected the unit’s time series of regression coefficients from Eq. 3 (β_*Reward*_ and β_*Delay*_) into a space in which reward size and delay compose two orthogonal dimensions. For each neuron, vector time series were produced in this regression space for significant β values (*P*<0.05) to capture the moment-by-moment mixture of encodings (Fig. 3C). In this space, vector angles indicated how a neuron’s activity reflected the combined effects of reward size and delay, while vector length captured the strength of the combined encoding. To determine the predominant encoding of these two characteristics (angle and length of moment-by-moment vectors) for each neuron, we summed across the time-resolved vectors (e.g., red dashed lines in Fig. 3C). The angles (θ) of the resulting vector sums were used to identify patterns of activity consistent with, and those inconsistent with, encoding of a temporal discounting of reward value – that is, an encoding in which reward size and delay have opposing effects on firing rate (Fig. 4A). Some neurons exhibited primarily a positive encoding of reward combined with a negative encoding of delay (vector angles −90°<θ<0°; referred to as the ‘Discounting−’ pattern in Fig. 4A; see e.g., Fig. 3C *right*), while others modulated their activity in the converse pattern with a negative β_*Reward*_ and positive β_*Delay*_ value (90°<θ<180°; referred to as ‘Discounting+’ pattern; see e.g., Fig. 3C *middle*). Other neurons encoded reward size and delay in an additive fashion, inconsistent with a signal reflecting subjective value and referred to here as ‘Compounding+’ and ‘Compounding−’ patterns (Fig. 4A). Compounding signals like these are inconsistent with a temporal discounting of reward value and may instead be attributable to extraneous factors such as arousal or attentional engagement.

**Figure 4.**
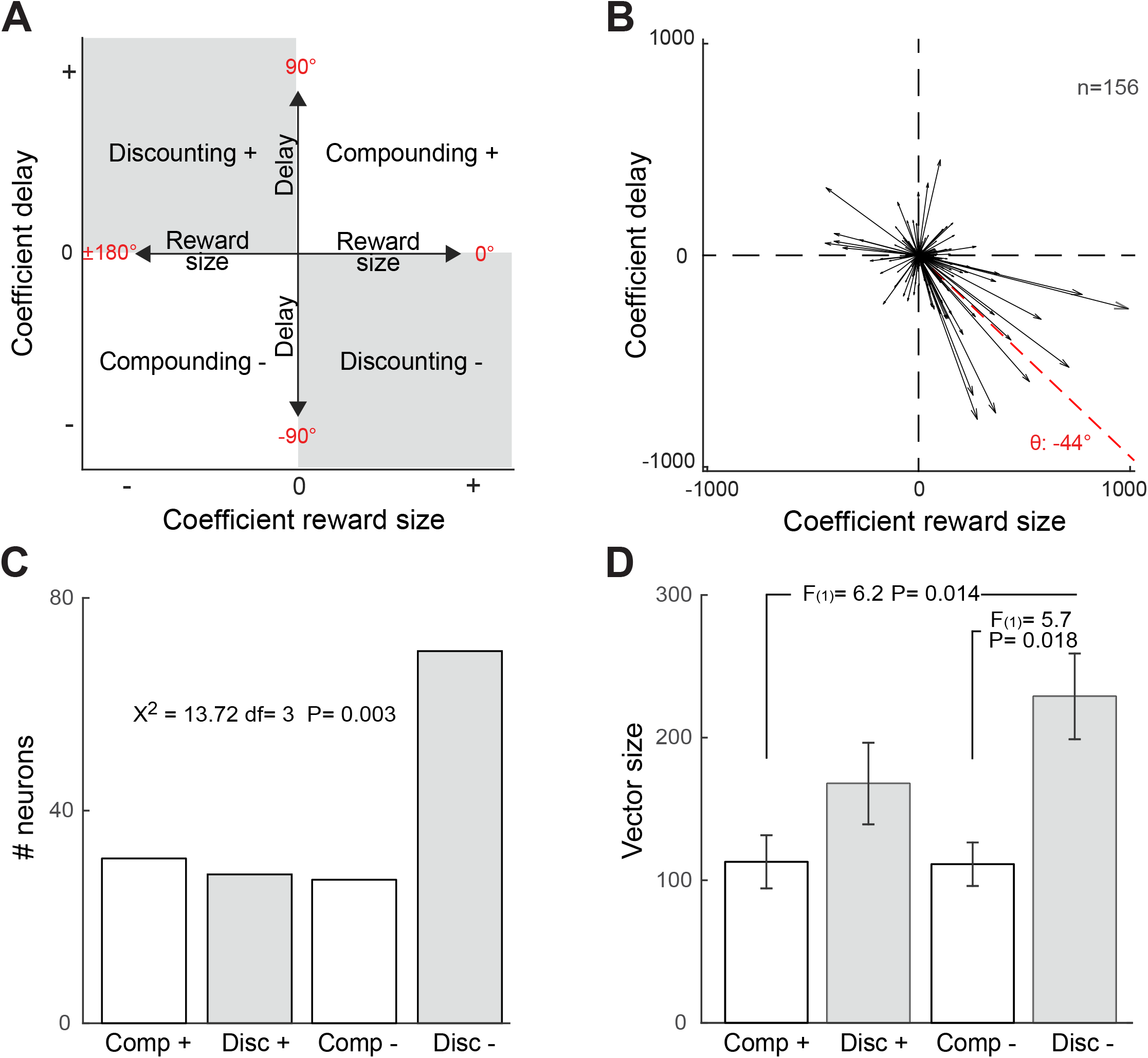
STN neurons exhibit signals consistent with temporal discounting. (**A**) Schematic depiction of the regression subspace composed of reward size and delay. Various patterns of neural encoding could be categorized depending on the angle (θ) of vectors: Discounting− (between −90 and 0°); Discounting+ (between 90 and 180°); Compounding+ (between 0 and 90°); Compounding− (between −180 and −90°). (**B**) Vectorial encoding of reward size and delay for all task-related neurons. Vector sums were calibrated by subtracting the mean 0 values of the pre-instruction epoch and then dividing by 2 SD of this control period. The red dashed line indicates the population vector. (**C**) Fractions of task-related neurons categorized as Discounting cells (Disc−, Disc+) or Compounding cells (Comp+, Comp−). (**D**) Vector sizes (mean ± SEM) were compared between different categories of task-related neurons (ANOVA).

At the population level, the neural encoding of reward size and delay parameters predominantly followed a Discounting pattern as evidenced by the fraction of neurons with Discounting− type encodings (χ^2^=13.72 *P*=0.003; Fig. 4B-C), and the longer mean vector length of Discounting− units (*F_(3,153)_*=4.06 *P*=0.008; Fig. 4D). Of the 156 task-related neurons, 70 (45%) increased firing rates as a function of reward size while they decreased according to the temporal delay to reward delivery (i.e, consistent with a Discounting− pattern). The remaining neurons were distributed equally across the other three encoding patterns. Vector lengths for neurons with a Discounting− firing pattern were longer, on average, than those for neurons with either type of Compounding pattern, while the vector length for neurons with a Discounting+ pattern fell in-between. Notably, the angle of the mean vector across all neurons (θ = −44°, Fig. 4B) showed that, despite the wide diversity of encoding patterns across individual neurons, the whole neural ensemble encoded information about reward size and delay in a pattern that was strongly consistent with a temporal discounting of value (i.e, Discounting−). We used a shuffled control procedure across directions of individual vectors to confirm that the length of the mean population vector was significantly greater than what would be expected by chance (*P*=0.004). Hence, the neural ensemble combined information related to reward size and delay into a coherent population-scale signal that reflected subjective value according to a Discounting− pattern.

Remarkably, the neurons categorized as Discounting− were located preferentially in the most posterior portion of the STN (2-tailed *t*-test, *t*_(229)_=3.37 *P*=0.0008; Fig. 5A-B). We did not observe a strict anatomic segregation of neurons with different encoding patterns, but rather an intermixed gradient of cell types along the antero-posterior axis. Discounting− neurons did not differ from other types of STN neurons with respect to tonic firing rate (*t*_(229)_=1.75 *P*>0.08; Fig. 5E) or action potential shape (*t*_(229)_<0.5 *P*>0.62; Fig. 5F-G). Aside from the posterior bias to their position in the STN, the neurons categorized as Discounting− were indistinguishable from the others.

**Figure 5.**
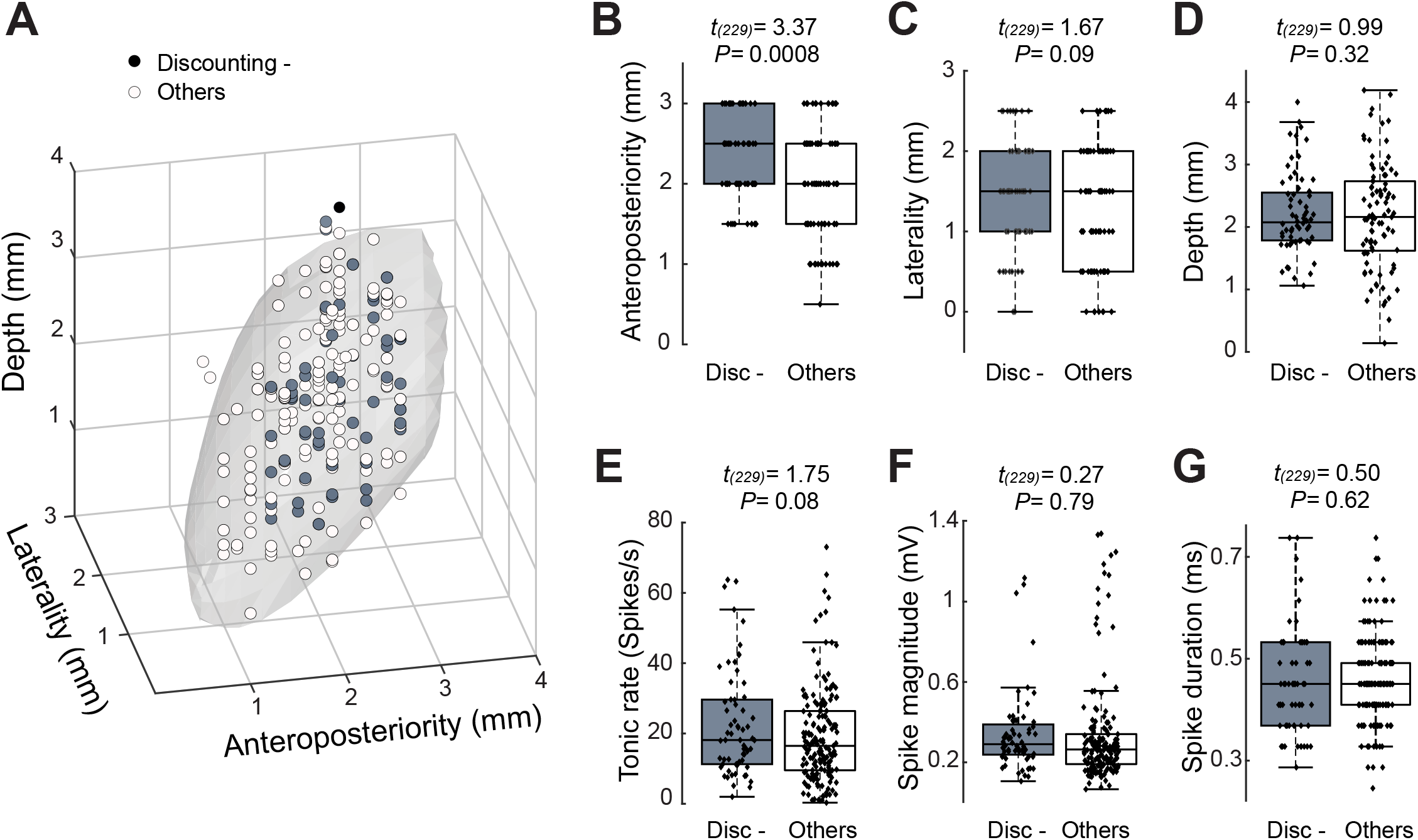
Topography of STN neurons categorized as Discounting−. (**A**) Three-dimensional plots of cell type distributions based on coordinates from the recording chamber. The template of STN (*gray surface*) is derived from the atlas Martin and Bowden (1996). (**B-D**) Comparisons of cell positions show that Discounting− neurons (Disc-) were located more posterior than other categories of cell type (*t*-test). (**E-G**) No differences were found in the tonic rate (**E**), spike magnitude (**F**), and spike duration (**G**) between Discounting− neurons and other cells. The central line of the box plots represents the median, the edges of the box show the interquartile range, and the edges of the whiskers show the full extent of the overall distributions.

### Dynamic encodings by the neural population ensemble

Using all recorded neurons (n=231), we then performed a population-based analysis to identify the principal representations of reward size and delay within the neural ensemble and how they changed dynamically over the course of a trial. After a projection of time series of regression coefficients (β_*Reward*_ and β_*Delay*_) into the orthogonal space composed of the reward size and delay, we used a principal component analysis (PCA) to identify the predominant patterns (i.e., the Principal Components, “PCs”) of encoding across the population. Given that the activity of individual neurons encoded diverse time-varying representations of task-relevant information (see e.g., Fig. 3), the PCA identified the patterns that accounted for the greatest variance within the neural population. In the resulting time series of eigenvectors, the angle of the eigenvectors indicated how reward size and delay parameters were integrated at the population level, while the length of eigenvectors captured the relative strength of the representations in the neural ensemble.

We identified the PCs that accounted for a significant faction of variance relative to that accounted for by a population of control PCAs (Fig. 6A-B). Surrogate control PCAs were computed by shuffling neural activity across trials before application of the PCA, thereby scrambling the relationship between task conditions and the neural activity. We found that the first four PCs from the real data exceeded the 95% confidence interval of variances accounted for by PCs from the surrogate control PCAs. In total, the first four PCs explained 75% of the total variance (Fig. 6A). These four PCs were distinct from each other with respect to the pattern represented (Fig. 6E) and how it evolved across time in a trial (Fig. 6D). We identified the timing of significant representations of reward size and delay in PCs by comparing the length of eigenvectors to the 95% confidence interval of those calculated from the surrogate control PCAs (Fig. 6F). Consistent with our single-unit analyses, the first principal component (PC1, which accounted for 45.4% of variance) corresponded to a temporal discounting of reward value (i.e. a signal that increased with increasing reward size and decreased with increasing delay; −90°<θ<0°). As evidenced by the eigenvector time series of PC1, the neural ensemble signaled a slightly delay-discounted reward value in a stable manner for approximately 2-s after the presentation of the instruction cue and then deviated progressively toward a more delay-influenced pattern during the waiting period immediately preceding reward delivery. Thus, reward size and delay were combined into a common neural ensemble signal that evolved dynamically across the trial period. This dynamic integration contrasted strongly with the signal captured by PC2 (second column of Fig. 6; accounting for 13% of variance) in which the reward and delay dimensions were represented independently of each other. Although the PC2 population signal was sensitive to both task parameters, discrete changes were detected consecutively, first for reward size (0°), in the 1-s after presentation of the instruction cue, and then delay to reward delivery (90°), during the 2-s immediately before reward delivery. Note that the population activity captured by PC2 increased with longer delays to reward. The separation of reward and delay encoding in PC2 to non-overlapping task time periods made this pattern inconsistent with an ensemble representation of subjective integrated value.

**Figure 6.**
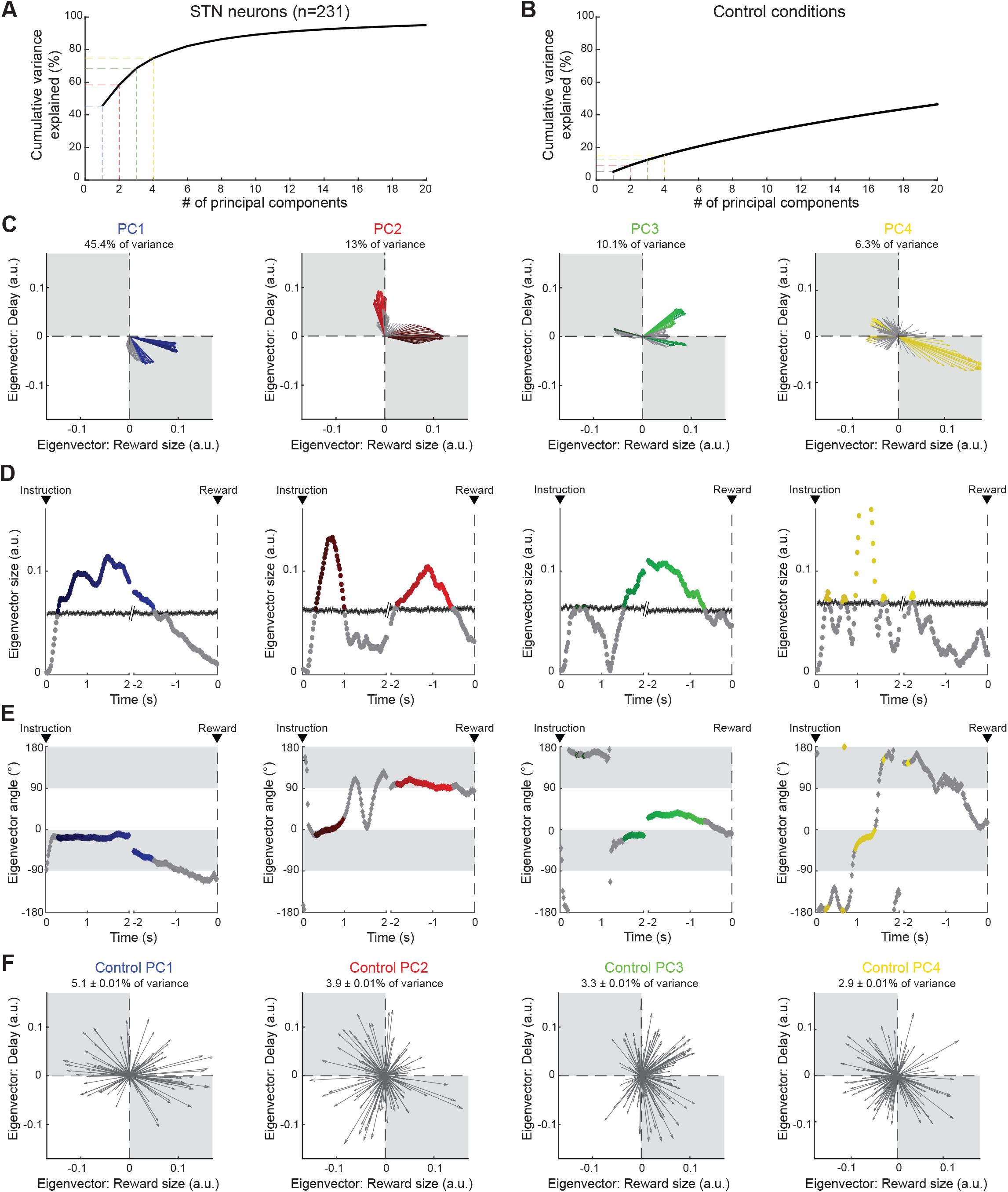
Neural population ensemble provides dynamic integration of reward size and delay. (**A**) Cumulative variance explained by PCA in the population of all recorded neurons. Dashed lines indicate the percentages of variance explained by the first four PCs: PC1 (*blue*), PC2 (*red*), PC3 (*green*) and PC4 (*yellow*). (**B**) Cumulative variance explained by a control shuffled procedure (data shuffled 1000 times). (**C**) Series of eigenvectors produced for the first four PCs. Eigenvectors capture the moment-bymoment signals in the subspace composed of reward size and delay. Colors indicated the eigenvectors with a significant length relative to those calculated from the surrogate control PCAs. (**D-E**) For each PC, eigenvector sizes captured the extent of the signals dynamically transmitted (**D**), while angles (−180 to 180°) indicated how the ensemble integrates reward size and delay (**E**). (**F**) Examples of eigenvectors produced by the control shuffled procedure. Percentages of variance explained by the first four PCs were significantly higher than chance (*P*<0.05).

The eigenvector properties for PC3 revealed a more complex population signal, which explained 10.1% of total variance. Between 1-2s after presentation of the instruction cue, PC3 approximated a short Discounting− type pattern of subjective value encoding (90°<θ<0°), after which, PC3 deviated to a robust Compounding+ pattern (0°<θ<90°) in which neural signals correlated positively with both the reward size and the time delay. The succession of these two phases suggests a switch in the functional role of task-relevant information when the waiting delay became longer in the post-instruction period. Finally, the signal extracted by PC4 explained 6.3% of variance in the ensemble and was characterized by a salient signal reflecting subjective values in a relative short manner at approximatively 1-s after instruction. Consistent with our single-unit analyses, PC4 corresponded also to a temporal discounting of reward value pointing steeply toward −21°.

### An anatomical gradient through STN

To test whether the predominant patterns of population encoding (i.e. PCs) described anatomical space in STN, we correlated the component scores of the neurons with their position along variable anatomical axes (45° rotations along x-y-z axes). Two anatomical axes correlated significantly with PC1 and PC4, i.e. the two components consistent with a processing of subjective value (Fig. 7). The scores for PC1 increased along the ventro-anterior to dorso-posterior axis in the STN (Spearman’s Rho=0.32 *P*<0.001; Figs. 7A, C), whereas the scores for PC4 increased strictly toward the most posterior portion of the STN (Rho=0.21 *P*=0.0014; Figs. 7B, D). Together, our data indicates that reward size and delay were primarily integrated into a subjective value along an antero-posterior gradient in the STN. No significant correlations were found in the spatial distribution of PC2 and PC3 scores (Rho<0.12 *P*>0.067), suggesting that these population signals were processed by neurons anatomically intermixed with others in the entire STN.

**Figure 7.**
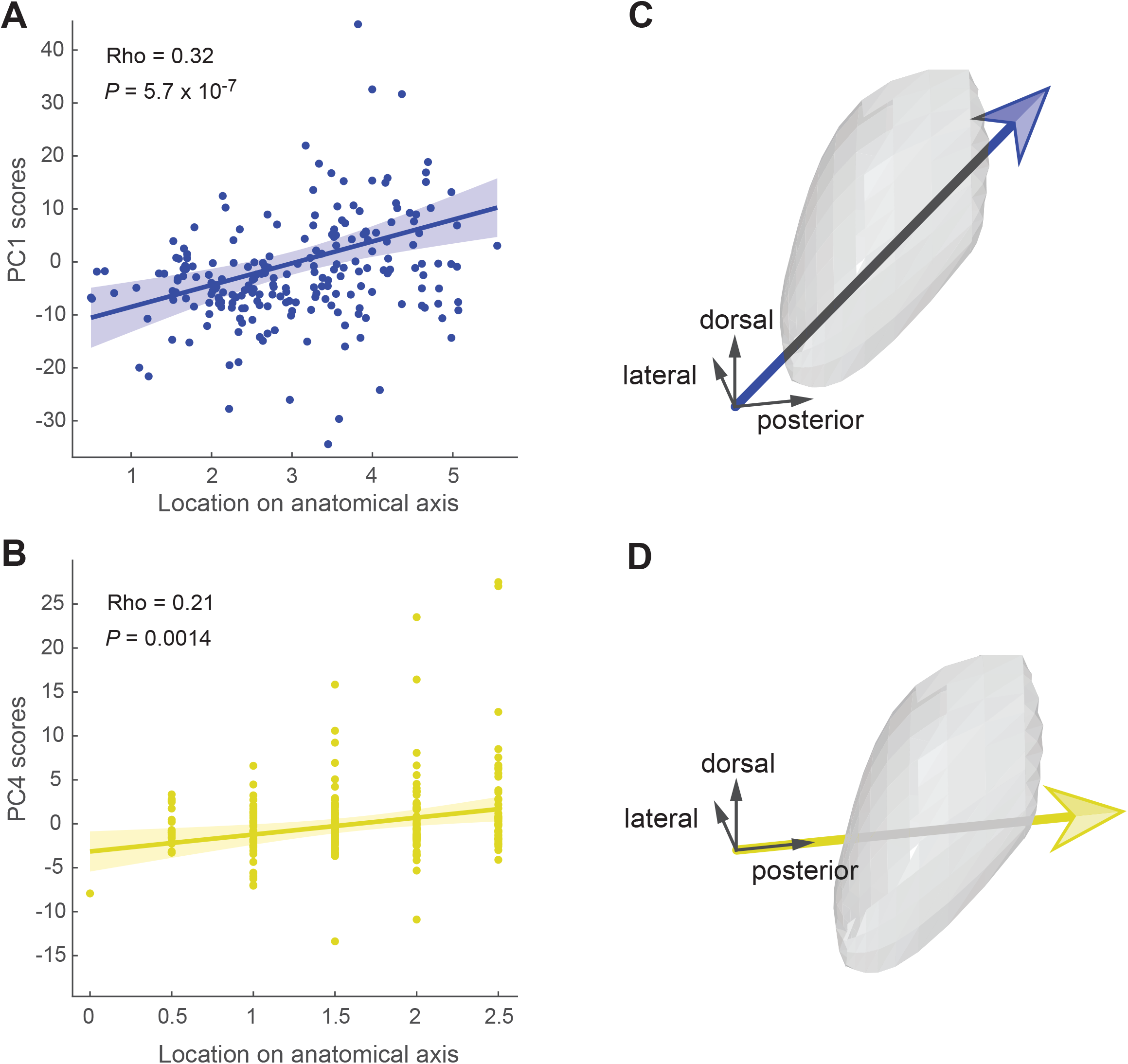
Signals vary along the antero-posterior axis of the STN. (**A-B**) Correlations between component scores [PC1 (*blue*) and PC4 (*yellow*)] of the neurons and their anatomical position. (**C-D**) Anatomical axis corresponding the most to PC1 (**C**) and PC4 (**D**). Arrows show direction of increasing component scores. No significant correlations were found in the spatial distribution of PC2 and PC3 scores (*P*>0.05).

## DISCUSSION

The present results reveal how STN neurons process temporally discounted value when a behavioral inhibition needs to be patiently sustained over time prior to delivery of a reward. At the single-neuron and population levels, we found a cost-benefit integration between the desirability of the expected reward and the imposed delay to delivery, with signals that dynamically combined both reward-related attributes to form a single integrated value estimate. The computation for such subjective value was increasingly observed along the antero-posterior axis of the STN, revealing that the most posterior-placed neurons are the most strongly involved in the representation of temporal discounting of value. These results expand our understanding concerning the involvement of STN in motivation and inhibitory control, by providing evidence for complex dynamical codings consistent with a continuous cost-benefit estimation promoting the goal pursuit and the willingness to bear the costs of time delays.

To determine if the STN conveys dynamic signals consistent with its predicted role in pursuing delayed gratification, we trained monkeys to perform a delayed reward task in which they were required to remain motionless for varying periods of time before the delivery of reward. We designed this task to investigate the ability of our animals to delay gratification and maintain a continuous self-control according to different levels of motivation. Previous studies have already tested monkeys’ behavior for delay maintenance (Evans et al., 2012; Evans and Beran, 2007; Freeman et al., 2012, 2009; Perdue et al., 2015; Szalda-Petree et al., 2004), but none to our knowledge have investigated the underlying neural mechanisms. Our analysis of animals’ performance confirmed that different levels of motivation or subjective value influenced their patience and willingness to sustain behavioral inhibition over time (Fig. 1). As evidenced by the more frequent rejection of trials offering small rewards and long delays, both cost-benefit parameters were integrated by monkeys to complete trials and finally obtain rewards. Consistent with previous findings (Fujimoto et al., 2019; Minamimoto et al., 2009), the likelihood of staying engaged in the trials for delayed rewards was well approximated by a model that calculates the subjective value with a hyperbolic delay discounting. This shows that monkeys estimated the cost-benefit conditions properly, and that active motivational processes were recruited during the time course of the post-instruction waiting period to maintain their behavioral inhibition.

In both human and non-human animal research, temporal discounting has been traditionally studied via the intertemporal choice task, which pits a small reward available sooner against a large reward available later (Frederick et al., 2002). As the delay to the large reward becomes longer, agents tend to start discounting the value of the large reward, biasing their preference toward the small reward available sooner. This choice behavior is considered impulsive and it referred to as a failure of self-control because it would be more economical to wait for the larger reward (Ainslie, 1975; Rachlin, 2004). To interpret individual choice behavior, the extent to which rewards are devalued over time have been preferentially inferred from hyperbolic discounting models (Mazur, 2001; Vanderveldt et al., 2016). Based on this approach, neuroimaging studies have identified a network of brain areas – involving the striatum, medial prefrontal cortex and posterior cingulate cortex – activation of which correlates with the subjective value of delayed rewards during decision-making processes (Kable and Glimcher, 2007; McClure et al., 2007, 2004; Peters and Büchel, 2009; Pine et al., 2009). Consistent with a coding of cost-benefit integration for delays to reward, BOLD activity in these regions increases as the expected amount of a reward increases and, inversely, decreases as a function of the imposed delay to reward. At the single neuron level, however, electrophysiological results have been more mixed concerning the integration of both reward-related attributes into a common currency value attribution (Roesch and Bryden, 2011). While some studies have found strict dissociable representations of both neural signals by different populations of neurons (Roesch et al., 2007, 2006; Roesch and Olson, 2005), two monkey studies have reported single-unit activities co-modulated by both reward size and delay (Cai et al., 2011; Kim et al., 2008). Our results confirm and extend to the STN these findings by showing that the cost and benefit dimensions of the task are represented in different ways in different sub-groups of neurons, either independently or in an integrated manner during the waiting period. Respectively, 15% and 14% of STN neurons were exclusively modulated by the expectation of reward size and delay without any co-modulation by the other parameter, maintaining distinct dynamic encodings split between these subsets of cells. Signals related to reward size were preferentially represented shortly after cue presentation, while signals related to delay cost occurred primarily later when animals endured the waiting period (Fig. 2). The fact we were able to dissociate in the STN these two types of neurons suggests that temporal discounting originates likely from different neural circuits than those that signal expected reward value. Alternatively, 38% of our STN neurons exhibited a dual coding of size and delay information, engendering a representation of both cost and benefit into a common dimension. Among those cells, the most prevalent activity pattern was characterized by antagonistic effects of reward magnitude and delay on their firing (Fig. 4). Specifically, STN neurons that fired more strongly for larger anticipated rewards tended to fire less strongly for longer delays (referred as Discounting−), consistent with the computation of temporally discounted value. Despite a high variability across individual neurons, we found that the whole neural ensemble sampled from the STN encoded and combined additively both reward size and delay into a strong population signal corresponding to a temporal discounting of reward value. As evidenced by the eigenvector time series of PC1, the integration of reward size and delay into this common signal was processed in a dynamic manner with time-variable representation of subjective values while animals remained immobile, waiting for the reward (Fig. 6). The longer the delay, the stronger was the encoding of waiting cost relative to the encoding of reward. Such dynamics appear consistent with the hypothesized role for STN in self-control and perseverance.

Unlike most studies of temporal discounting that focus on decision-related neural processes at just one time point in the task, we investigated activity during the post-instruction delay period, during which animals were required to exert sustained commitment to an action with a continuous behavioral inhibition. It is quite likely that distinct components of self-control are measured during those two time periods (Addessi et al., 2013). Our approach offers the opportunity to investigate the dynamic processes engaged during the post-instruction period to support an animal’s ability to delay gratification. An important avenue for future research will be to determine how STN signals, such as those described here, change when animals run out of patience and finally decide to stop waiting. To do this, however, smaller reward sizes and longer delays might be used to promote more escape behaviors during delay interval. Rejection rates were relatively low in the present version of our task. Given current models where STN is thought to prevent hasty decisions by elevating its activity to pause cortical commands via pallido-thalamocortical circuits (Cavanagh et al., 2011; Frank, 2006; Mansfield et al., 2011), signals underlying a steeper temporal discounting may be observed shortly before behavioral disinhibition if this nucleus effectively drives this type of inhibitory function. In addition, further studies relying on simultaneous multisite recordings are still necessary to clarify the neural origins of the information transmitted by the signals identified here. Although our results show that STN neurons process a dual dynamic coding of size and delay information, it remains undetermined whether the integration between these two reward-related attributes occurs primarily within this nucleus or upstream in other brain areas such as the dorsolateral prefrontal cortex (Kim et al., 2008) and the anterior striatum (Cai et al., 2011), for instance.

A growing number of animal studies have examined the STN involvement in motivational functions using single-unit recordings. Neurons with phasic responses evoked by reward-predictive cues and the reward itself have been reported in different species and tasks (Breysse et al., 2015; Darbaky et al., 2005; Espinosa-Parrilla et al., 2013; Lardeux et al., 2013, 2009; Matsumura et al., 1992; Nougaret et al., 2022; Teagarden and Rebec, 2007). However, it has been unclear how this reward processing relates to the known organization of STN into anatomically and functionally distinct territories (Alexander et al., 1990; Nambu et al., 2002; Parent and Hazrati, 1995). The STN receives topographically-organized inputs from most regions of the frontal cortex (Haynes and Haber, 2013; Nambu et al., 2002), the pallidum (Karachi et al., 2005; Shink et al., 1996), and the parafascicular nucleus of the thalamus (Sadikot et al., 1992). Together, tracing studies have indicated that the posterior-dorsal-lateral STN is interconnected with circuits devoted to sensorimotor function, whereas associative- and limbic-related territories are found in progressively more anterior, ventral and medial regions of the STN (Haynes and Haber, 2013; Mettler and Stern, 1962; Monakow et al., 1978; Shink et al., 1996). Based on this, a widely accepted tripartite model divides STN into segregated motor, associative, and limbic regions that are thought to play distinct functional roles in motor control, cognition, and emotion (Parent and Hazrati, 1995). Surprisingly, in previous primate studies, reward-responsive neurons were found to be scattered throughout all parts of the STN (Darbaky et al., 2005; Espinosa-Parrilla et al., 2013; Nougaret et al., 2022), rather than showing a preferential location in the anterior portion which is held to be the zone with strongest connectivity to limbic structures (Haynes and Haber, 2013; Karachi et al., 2005). This lack of anatomical specificity has been confirmed in human recordings for which reward-modulated neurons were also identified in sensorimotor regions of the STN (Justin Rossi et al., 2017; Sieger et al., 2015). To date, this paradoxical observation was attributed to a bias in data collection and the inherent observational bias introduced in data collected as part of DBS implantation surgeries due to the posterior-dorsal-lateral location of the target in those surgeries. By accumulating 231 neurons distributed throughout all portions of the STN, our study provides a more complete survey of this nucleus for regional variations in the representation of reward-related information. At the single-neuron and population levels (Figs. 5, 7), we found that the valuation of delayed rewards was represented preferentially in the most posterior portion of the STN, thereby challenging predictions from classical tripartite models. While earlier studies suggested segregated functional subdivisions, now more recent evidence points toward overlapping territories (Alkemade and Forstmann, 2014; Emmi et al., 2020). Consistent with a functional convergence within this structure, our results show that motivational signals were conveyed by neurons located in areas traditionally identified as part of the sensorimotor circuits. Because muscle activities were not altered by reward contingencies during the task (Figs. 1H-I), we assume that these dynamic reward-related signals were primarily processed by cognitive circuits that monitor and manage goal achievement by translating motivational drives into behavioral perseverance. This role in self-control extends our understanding concerning the STN involvement in controlling impulsive behaviors (Aron et al., 2016; Jahanshahi et al., 2015; Zavala et al., 2015) and the willingness to work for food (Baunez et al., 2005, 2002). By providing evidence for dynamic cost-benefit valuation by STN, our results imply that this structure promotes the pursuit of motivated behavior by computing which costs are acceptable for a reward at stake. Further research is needed to determine whether the neural signals identified here causally drive animals’ behavior or rather just participate to evaluate the current situation.

Impatience for reward and lack of perseverance are major facets of many psychiatric disorders. Numerous clinical studies have shown that these maladaptive behaviors are characterized by a disruption in the ability to weigh appropriately the amount of reward against the cost of delay (Kirby et al., 1999; Madden et al., 1997; Mitchell, 1999; Reynolds, 2006; Vuchinich and Simpson, 1998). For instance, patients with self-control issues such as in gambling disorder (Alessi and Petry, 2003), drug addiction (Kirby and Petry, 2004; Madden et al., 1997; Washio et al., 2011), schizophrenia (Ahn et al., 2011; Heerey et al., 2007; MacKillop and Tidey, 2011), depression (Dombrovski et al., 2012; Pulcu et al., 2014; Takahashi et al., 2008), mania (Mason et al., 2012), attention deficit hyperactivity disorder (Barkley et al., 2001; Scheres et al., 2010; Tripp and Alsop, 2001), and anxiety disorder (Rounds et al., 2007) have higher delay discounting rates than normal subjects. Our results merge existing lines of evidence that implicate the STN in motivation and inhibitory control, positioning this structure as a potential hub to regulate aberrant reward processing and the capability to postpone. This view agrees with beneficial effects of DBS-STN on drug addiction reported in recent animal studies (Pelloux et al., 2018; Rouaud et al., 2010; Wade et al., 2017), and with the battery of psychiatric side-effects observed in parkinsonian patients after electrode implantation within the STN (Castrioto et al., 2014). However, it remains an open question how the STN contributes to distort defective valuation processes in terms of cost-benefit trade-off in these pathological states.

## MATERIALS AND METHODS

### Animals

Two rhesus monkeys (monkey C, 8 kg, male; and monkey H, 6 kg, female) were used in this study. Procedures were approved by the Institutional Animal Care and Use Committee of the University of Pittsburgh (protocol number: 12111162) and complied with the Public Health Service Policy on the humane care and use of laboratory animals (amended 2002). When animals were not in active use, they were housed in individual primate cages in an air-conditioned room where water was always available. The monkeys’ access to food was regulated to increase their motivation to perform the task. Throughout the study, the animals were monitored daily by an animal research technician or veterinary technician for evidence of disease or injury and body weight was documented weekly. If a body weight <90% of baseline was observed, the food regulation was stopped.

### Behavioral task

Monkeys were trained to perform the delayed reward task with the left arm using a torquable exoskeleton (KINARM, BKIN Technologies, Kingston, Ontario, Canada). This device had hinge joints aligned with the monkey’s shoulder and elbow and allowed the animal to make arm movements in the horizontal plane. Visual cues and cursor feedback of hand position were presented in the horizontal plane of hand movements by a virtual-reality system. A detailed description of the apparatus can be found in our previous studies (Pasquereau and Turner, 2015, 2013).

In our delayed reward task (Fig. 1A), the monkey was required to align the cursor on a visual target (radius, 1.8 cm) and to maintain this position for varying periods of time before delivery of the food reward. In total, six combinations of reward size and waiting delay were used. A trial began when a gray-filled target appeared (the same location for all trials) and the animal made the appropriate joint movements to place the cursor in this circle. Maintenance of the cursor within the target required the animal to actively stabilize the posture of both shoulder and elbow joints in the horizontal plane. While the monkey maintained its arm position, an instruction cue was displayed over the gray-filled target for 0.5 s. After a variable interval (1.2-2.8 s), the gray fill disappeared from the circle, cueing the animal to remain motionless during an additional delay until the reward delivery. For the instruction cues, cue colors indicated the size of reward (1, 2, and 3 drops of food) and symbols indicated the delay duration that the animal would have to wait between the gray fill disappearance and reward delivery [short delay (1.8-2.3s) and long delay (3.5-4s)]. Cue colors were calibrated to have the same physical brightness (~30 cd/m2). The six unique cue types (3 reward sizes x 2 delay ranges) were presented in pseudo-random order across trials with equal probability. At the end of each successful trial, food reward was delivered via a sipper tube attached to a computer-controlled peristaltic pump (1 drop = ~0.5 ml, puree of fresh fruits and protein biscuits). The trials were separated by 1.5–2.5 s intertrial intervals, during which the screen was black. Failures to maintain the cursor in the start position (radius = 1.8 cm) between instruction cue and reward delivery were counted as errors. After an error, the rejected trial was aborted and a blank scream appeared (1s), followed by an intertrial interval.

To confirm the ability of our monkeys to evaluate appropriately the different instruction cues, we also trained them to perform a decision task in which they were allowed to choose freely between two alternate reward size/delay combinations. The overall structure of this task was similar to the delayed reward task except two instruction cues were presented simultaneously. The location of the cues alternated randomly between left and right sides. The animal chose its preferential option by positioning and maintaining the cursor on one of the targets. The pair of cues presented on a single trial differed only along one dimension, presenting either a reward-based decision (different colors but the same symbol) or a delay-based decision (different symbols but the same color).

Before the start of data collection, we trained the monkeys to perform these two behavioral tasks for more than 6 months. Neuronal data were not collected during performance of the decision task.

### Surgery

After reaching asymptotic task performance, animals were prepared surgically for recording using aseptic surgery under Isoflurane inhalation anesthesia. An MRI-compatible plastic chamber (custom-machined PEEK, 28 × 20 mm) was implanted with stereotaxic guidance over a burr hole allowing access to the STN. The chamber was positioned in the parasagittal plane with an anterior-to-posterior angle of 20°. The chamber was fixed to the skull with titanium screws and dental acrylic. A titanium head holder was embedded in the acrylic to allow fixation of the head during recording sessions. Prophylactic antibiotics and analgesics were administered post-surgically.

### Localization of the recording site

The anatomical location of the STN and proper positioning of the recording chamber to access it were estimated from structural MRI scans (Siemens 3 T Allegra Scanner, voxel size of 0.6 mm). An interactive 3D software system (Cicerone) was used to visualize MRI images, define the target location and predict trajectories for microelectrode penetrations (Miocinovic et al., 2007). Electrophysiological mapping was performed with penetrations spaced 1 mm apart. The boundaries of brain structures were identified based on standard criteria including relative location, neuronal spike shape, firing pattern, and responsiveness to behavioral events (e.g. movement, reward). By aligning microelectrode mapping results (electrophysiologically characterized X-Y-Z locations) with structural MRI images and high resolution 3-D templates of individual nuclei derived from an atlas (Martin and Bowden, 1996), we were able to gauge the accuracy of individual microelectrode penetrations and determine chamber coordinates for the STN.

### Recording and data acquisition

During recording sessions, a glass-coated tungsten microelectrode (impedance: 0.7–1 MOhm measured at 1000 Hz) was advanced into the target nucleus using a hydraulic manipulator (MO-95, Narishige). Neuronal signals were amplified with a gain of 10K, bandpass filtered (0.3–10 kHz) and continuously sampled at 25 KHz (RZ2, Tucker-Davis Technologies, Alachua FL). Individual spikes were sorted using Plexon off-line sorting software (Plexon Inc., Dallas TX). The timing of detected spikes and of relevant task events was sampled digitally at 1 kHz. Horizontal and vertical components of eye position were recorded using an infrared camera system (240 Hz; ETL-200, ISCAN, Woburn, MA). For electromyographic recordings (EMG, in monkey H only), pairs of Teflon-insulated multi-stranded stainless-steel wires were implanted into multiple muscles during 12 training sessions. EMG signals were differentially amplified (gain = 10 K), band-pass filtered (200 Hz to 5 kHz), rectified and then low-pass filtered (100 Hz).

### Analysis of behavioral data

We analyzed the way the animals performed the delayed reward task to test whether the behavior varied according to the levels of reward and delay. As the monkey maintained its arm position for varying periods of time before to obtain rewards, the sum of errors per session was the main variable of interest. Rejection rates were calculated by dividing the number of errors by the total number of trials for each task condition. Two-way ANOVAs were used to test these rejections for interacting effects of reward size (1, 2, or 3 drops) and delay (short or long delay duration). Modulation in rejection rates reflected the monkey’s motivation to stay engaged in the task and to fully complete the different types of trials by maintaining the correct arm position. In the present study, the motivational or subjective value linked to each task condition was estimated by integrating the forthcoming reward size and the delay discounting. The subjective value of delayed reward is commonly formulated as a hyperbolic discounting model (Green and Myerson, 2004; Mazur, 1987) as follows:

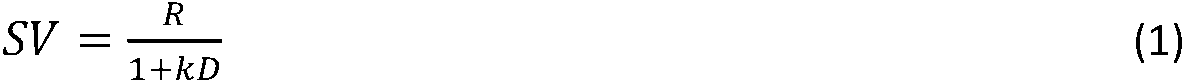

where *SV* is the subjective value (i.e. the temporally discounted value), *R* is the reward size, *k* is a discount factor that reflects an individual animal’s sensitivity to the waiting cost, and *D* is the delay to the reward. In previous studies (Hori et al., 2021; Kobayashi and Schultz, 2008; Minamimoto et al., 2009), this hyperbolic discounting model has been consistently shown to be better than an exponential function to fit to the monkeys’ behavior. Because the number of errors in task performance is inversely related to the subjective value (Fujimoto et al., 2019; Minamimoto et al., 2009), we have inferred the subjective value in each monkey by fitting the average rejection rate to the following model:

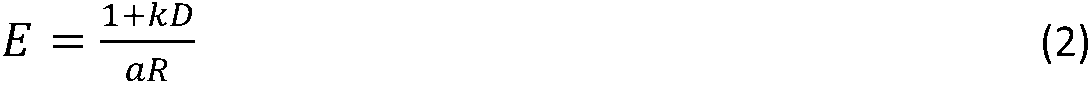

where *E* is the rejection rate, *R* is the reward size, *D* is the delay, *k* is a discount factor, and *a* is a monkey-specific free parameter. We estimated the best pair of free parameters (*k* and *a*) with the Matlab function “fminsearch” that provided the maximum-likelihood fit. Goodness of fit was evaluated by the coefficient of determination (R^2^).

As monkeys were not required to control their gaze while performing the task, we tested whether eye positions varied according to the levels of reward and delay. For this analysis, we combined horizontal and vertical components to obtain tangential coordinates, and the potential interaction with task parameters was examined using a two-way ANOVA combined with a sliding window procedure (200-ms test window stepped in 20-ms). The threshold for significance (*P* < 0.05, corrected for n-time bins) was validated by calculating the likelihood of Type 1 (false positive) errors during the pre-instruction control epoch (i.e., when the animal cannot anticipate the upcoming reward and delay). The same statistical method was used to analyze EMGs, with series of two-way ANOVA performed to test for effects of reward size and delay. All of the data analyses were performed using custom scripts in the MATLAB environment (The MathWorks, MA).

### Neuronal data analysis

Neuronal recordings were accepted for analysis based on electrode location, recording quality (signal/noise ratio of >3 SD) and duration (>120 trials). The width of action potential waveforms was calculated as the interval from the beginning of the first negative inflection (>2 SD) to the subsequent positive peak, and the magnitude of the biphasic spike waveforms was measured between maxima and minima. Trials with errors were excluded from the analysis of neuronal data. For different task events (i.e., instruction cue and reward delivery), continuous neuronal activation functions [spike density functions (SDFs)] were generated by convolving each discriminated action potential with a Gaussian kernel (20-ms variance). The influence of task parameters on the activity of STN neurons was investigated across the full trial duration: from the epoch preceding the instruction cue (−1 to +2 s) to the time period of the reward delivery (−2 to +1 s). Mean peri-event SDFs (averaged across trials) for each of the reward and delay conditions were constructed. A neuron’s baseline firing rate was calculated as the mean of the SDFs across the 1 s epoch preceding cue instruction. The Fano factor, defined as the variance-to-mean ratio of firing rates, was used to measure the variability of neuronal activities across the six trial conditions. For each single-unit activity, we tested for effects of reward size and delay using two-way ANOVAs combined with a sliding window procedure (200-ms test window stepped in 20-ms). Specifically, we extracted single-trial spike counts from a series of 200-ms windows and we investigated for each step whether the neuronal activity was influenced by 1) reward size, 2) delay to reward, or 3) both task parameters. The ANOVA identified any interacting effects of reward size and delay. The threshold for significance (*P* < 0.05, corrected for n-time bins) was validated by calculating the likelihood of Type 1 (false-positive) errors in a parallel analysis of activity extracted from the 1 s of the pre-instruction control epoch. A neuron was judged to be task-related if it generated a significant encoding of the reward size and/or delay for a least one-time bin from instruction cue to reward onset.

To test how individual task-related neurons dynamically encoded the forthcoming reward and the delay discounting through trials, we used time-resolved multiple linear regressions. We simultaneously tested whether trial-to-trial neuronal activity was modulated by the size of reward (*R*), the delay duration (*D*), and measures of eye movements made in the task [tangential position (*P*) and tangential velocity (*V*)] to control for possible relationships of STN activity with gaze control. For each task-related neuron, we counted spikes (*SC*) trial-by-trial within a 200-ms test window that was stepped in 20-ms increments relative to task events (3 s around instruction and reward onset). For each bin, we applied the following model:

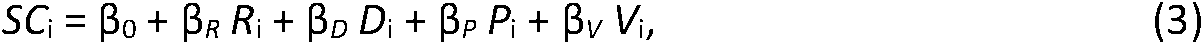

where all regressors for the i^th^ trial were normalized to obtain standardized regression coefficients (Z-scored in standard deviation units). The β coefficients were estimated using the ‘glmfit’ function in Matlab. The threshold for significance of individual β values was determined from the set of coefficients calculated within the 1 s of the pre-instruction control epoch (2-tailed *t*-test, *P* < 0.05). In addition, we estimated the relative efficiency of this model (as expressed by Eq. 3) by comparing the goodness-of-fit with another model where the neural activity was modulated by the subjective value (*SV*) calculated from the animal’s behavior (as expressed by Eq. 1). In this alternative model, spike counts were fitted as follows:

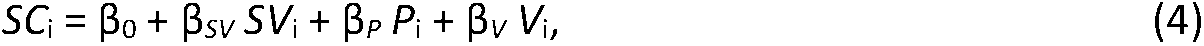

Models were compared based on Akaike information criterion (AIC), a measure of goodness-of-fit that penalizes models possessing more parameters (Akaike, 1974).

To characterize how individual neurons integrate dynamically task parameters, time series of regression coefficients of Eq. 3 (β_*R*_ and β_*D*_) were projected into an orthogonal space where reward size and delay composed the two dimensions (axes) of interest. In this regression space, we used significant points (pairs of β_*R*_-β_*D*_) to produce vector time series originating from the control value of the pre-instruction epoch (averaged across time bins). Vectors were generated with significant regression coefficients (2-tailed *t*-test, *P* < 0.05) calculated from instruction cue to reward onset. In this context, vector angles (−180 to 180°) indicated how the neural activity encodes and combines reward size and delay, while vector sizes captured the extent of the signals transmitted. Considering these two characteristics (direction and length of vectors), we summed all time-resolved vectors to identify the predominant encoding of task parameters for each neuron analyzed. Various patterns of neural encoding could be categorized from the angle (θ) of the resultant vector sum. For instance, θ indicated whether the neural activity was correlated (positive coding) or anticorrelated (negative coding) with the reward size, the delay to reward, or both. 0° (positive β_*R*_ values) indicated an exclusive positive reward size coding, ±180° (negative β_*R*_ values) indicated an exclusive negative reward size coding, 90° (positive β_*D*_ values) indicated an exclusive positive delay coding, and −90° (negative β_*D*_ values) indicated an exclusive negative delay coding. θ between −90 and 0°, or between 90 and 180°, indicated a coding of both reward size and delay consistent with a temporally discounted value where both benefit and cost parameters modulate the subjective representation in an opposite way (i.e, positive β_*R*_ with negative β_*D*_, or vice versa). θ between 0 and 90°, or between −180 and −90°, indicated a coding of reward size and delay where both parameters are integrated in a compound signal inconsistent with a temporally discounted value (i.e, positive β_*R*_ and β_*D*_, or negative β_*R*_ and β_*D*_). For population-based figures (Fig. 4), vectors were standardized between neurons by subtracting the mean values of the pre-instruction epoch and then dividing by 2 SD of this control period. The population vector was calculated by vectorially summing standardized cell vectors.

Using all recorded neurons, we then performed an alternative population-based analysis to characterize the predominant patterns of neural encoding in the STN. We used a principal component analysis (PCA) to identify patterns of encoding (dimensions) of the neuronal ensemble in the orthogonal space composed of the reward size and delay. Our procedure was analogous to the analyses performed by Yamada et al., 2021. A two-dimensional data matrix *X* of size *N*_(neuron)_ × *N*_(*C*×*T*)_ was prepared with regression coefficients of Eq. 3 (series of β_*R*_ and β_*D*_), in which rows corresponded to the total number of neurons and columns corresponded to the number of conditions (*C*, reward size and delay) multiplied by the number of time bins (*T*). A series of eigenvectors was obtained by applying PCA once to the data matrix *X*. In our analysis, the eigenvectors represent vectors at different time bins in the orthogonal space composed of reward size and delay. Principal components (PC) were estimated using the ‘pca’ function in Matlab. As for the cell vectors, vector angles (−180 to 180°) indicated how the ensemble encodes and combines reward size and delay, while vector sizes captured the extent of the signals dynamically transmitted. Adequate performance of PCA was estimated with the percentages of variance explained by PCs. To test whether the PCA performance was significant, we constructed a surrogate control population of PCs in which the neural activity (*SC*) was shuffled across trials before application of Eq. 3. Consequently, in the surrogate data the linear projection of neural activity into the regression subspace was randomized, eliminating any coherent modulation of activity with task parameters (reward size and delay) in the matrix *X*. We built a surrogate control population of PCs by repeating the shuffling procedure 1000 times and then compared percentages of variance explained by actual PCs against the 95% confidence interval of variances accounted for by the population of surrogate PCs. An actual percentage of variance estimated by PCA was deemed significant if its value fell outside of the 95% confidence interval of those for the surrogate. In addition, we identified the timing of significant representations of reward size and delay in PCs by comparing the length of eigenvectors to the 95% confidence interval of those calculated from the surrogate control PCA.

To test how PCs mapped onto anatomical axes in STN, we correlated the component scores of the neurons with their location (x-y-z coordinates). We performed this in standard stereotaxic space and also for a rotated set of anatomical axes (45° rotations along x-y-z axes). The rotated axis that best correlated with the scores of a component (maximal Spearman’s Rho) was identified as the optimal axes for explaining that components variance.

## SUPPLEMENTAL FIGURE LEGEND

**Figure 1 - supplement 1.**
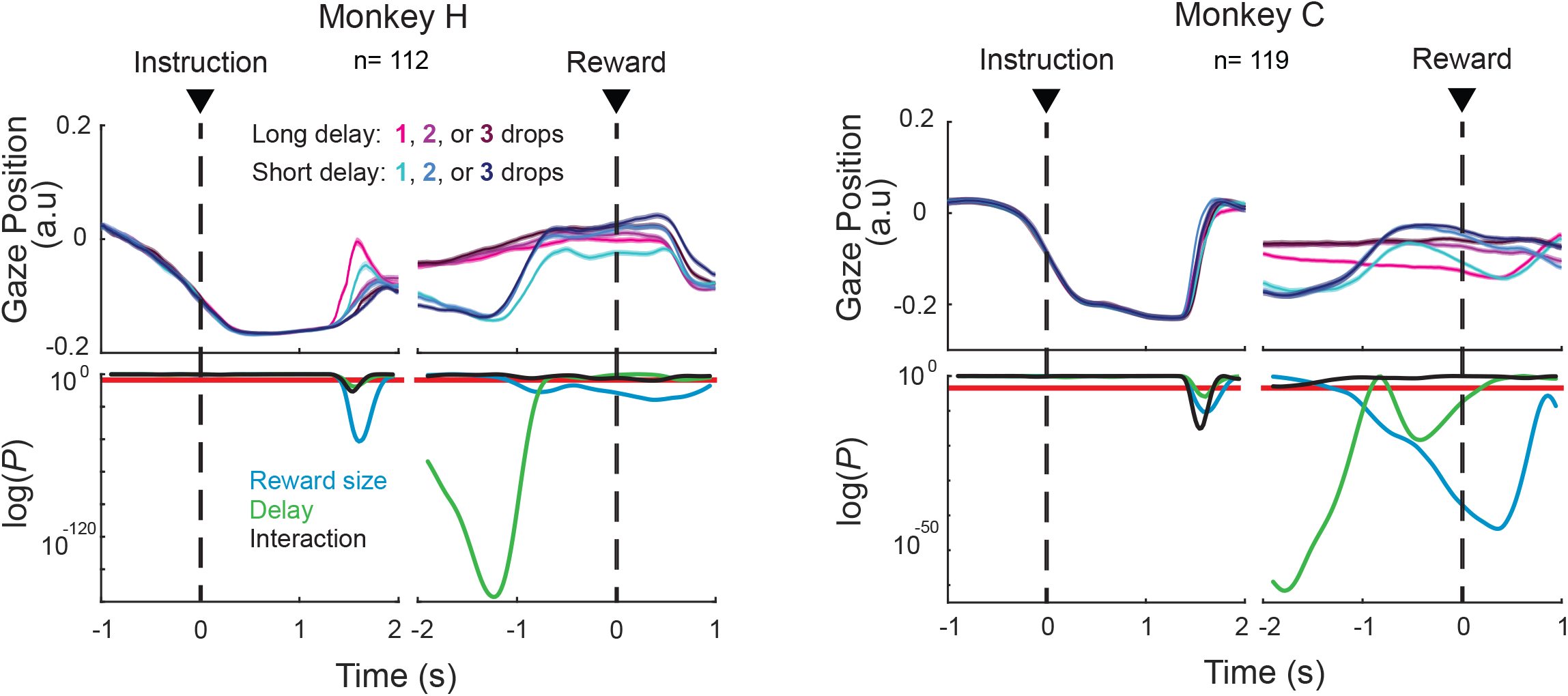
Eye positions varied according to the levels of reward and delay. For each task condition, horizontal and vertical components were combined to obtain tangential coordinates (averaged across sessions; ± SEM). Potential interaction with task parameters was tested using a series of two-way ANOVAs (Reward size × Delay). Red lines indicate the statistical threshold (*P*<0.05 corrected for 143-time bins). This threshold for significance was validated by calculating the likelihood of Type 1 (false positive) errors during the pre-instruction control epoch (i.e., when the animal cannot anticipate the upcoming reward and delay).

